# Spatial variability of shade and its effects on cocoa tree productivity in agroforestry systems

**DOI:** 10.1101/2025.11.06.687074

**Authors:** Guei Stéphane Hubert, Sanogo Souleymane, M’bo Kacou Alban Antoine, Thomas Wibaux, Chérif Mamadou, Patrick Jagoret

## Abstract

The impact of intra-plot multi shading on cocoa tree productivity is a widely exploited aspect of agroforestry practice. This study, conducted on 1,000 m2 experimental plots at three agroclimatic sites in Côte d’Ivoire (Azaguié, Divo, Soubré), mapped the spatial heterogeneity of shade provision using light intensity measurements and then calculated cocoa productivity over two years of data collection. A geostatic approach (ordinary kriging) was used to analyze the shade in agroforestry plots, which revealed significant heterogeneity. This heterogeneity was primarily attributed to the phenological diversity of the companion trees. The analysis revealed that low shading levels (approximately 30%) emerged as thresholds where cocoa productivity was highest. However, it was observed that elevated shading levels, exceeding 60%, resulted in a significant decline in productivity. Among the companion trees, *Ceiba pentandra, Eleais guineensis*, and *Persea americana* offer light to moderate shade. Conversely, certain trees that provide heavy shade, such as *Mangifera indica*, are identified as less beneficial because they significantly reduce cocoa productivity.

## 1. Introduction

Global climate projections highlight the growing threats to agricultural systems, particularly in tropical regions, where rising temperatures, rainfall variability, and prolonged droughts seriously compromise crop sustainability. These pressures exacerbate the negative effects of intensive agriculture and monoculture, which remain among the main drivers of deforestation, biodiversity loss, and soil degradation. In this context, adopting “climate-smart” agricultural practices is vital to sustainably balance productivity, resilience and conservation [1,2]. The cacao tree (*Theobroma cacao* L.), a species native to forest undergrowth [3,4], is particularly vulnerable to global warming, especially in monoculture, where its increased exposure to solar radiation intensifies thermal and water stress [5,7]. Recent projections indicate a substantial decline in productivity and a reduction in suitable areas for cocoa cultivation should no adaptation measures be implemented [8–10]. In the context of these challenges, agroforestry emerges as a potentially viable solution. The integration of shade trees into cocoa plantations plays a central role in regulating the microclimate, reducing heat and water stress, conserving soil moisture, and even controlling pests [11,12]. However, it is well established that the effectiveness of these systems depends largely on the species associated with them. Their morphological and architectural characteristics determine the amount and spatial distribution of the shade provided [13,14]. In this context, [15] highlight the importance of quantifying the effects of tree cover on the microclimate in greater detail. Several studies have confirmed that the architectural characteristics of shade trees directly influence the performance of cacao trees [16], their physiology depending on the distance between them and these trees [17], as well as the incidence of major pests such as mirids and brown rot [18]. However, gaps remain, as most studies focus on the average effects of shade or on specific species, without considering the high spatial variability of shade at the intra-plot level. This heterogeneity, known as “multishading” [19], generates contrasting light regimes that can differentially influence the growth, resilience, and productivity of cocoa trees within the same orchard. This dimension has been little explored, which is a gap in research on the optimization of agroforestry systems. Recent methodological advances, such as terrestrial laser scanning (TLS) and canopy hemispherical photography combined with machine learning or light mapping using the geospatial kriging method, now offer tools to analyze this spatial complexity in detail [20]. Understanding shade distribution, not just its average, is crucial for production. This study uses a geospatial approach to analyze how multiple shading affects cocoa production variability within plots.

## 2. Materials and methods

### 2.1. Study sites

The study was conducted on 1000 m^2^ experimental cocoa plots within traditional agroforestry farms at three locations in Côte d’Ivoire: Azaguie (5°37′ N, 4°05′ W) in a southern forest–savanna transition zone; Divo (5°50′ N, 5°21′ W) in an intermediate sub-equatorial zone; and Soubre (5°46′ N, 6°36′ W) in a humid forest zone in the southwest. These sites, among the country’s main cocoa-producing areas, have distinct agroecological conditions. Azaguie features a tropical savanna climate with approximately 1466 mm of annual rainfall and a mean temperature of 26.7 °C. Divo has a subequatorial climate (approx. 1200 mm annual rainfall, 26.5 °C average temperature). In contrast, Soubre experiences a humid tropical climate (approx. 1069 mm annual rainfall, 26.1 °C on average). These climatic variations created diverse environmental conditions, enabling a comprehensive assessment of the relationship between shade and cocoa yield (Fig 1) [21,22].

**Fig 1.**
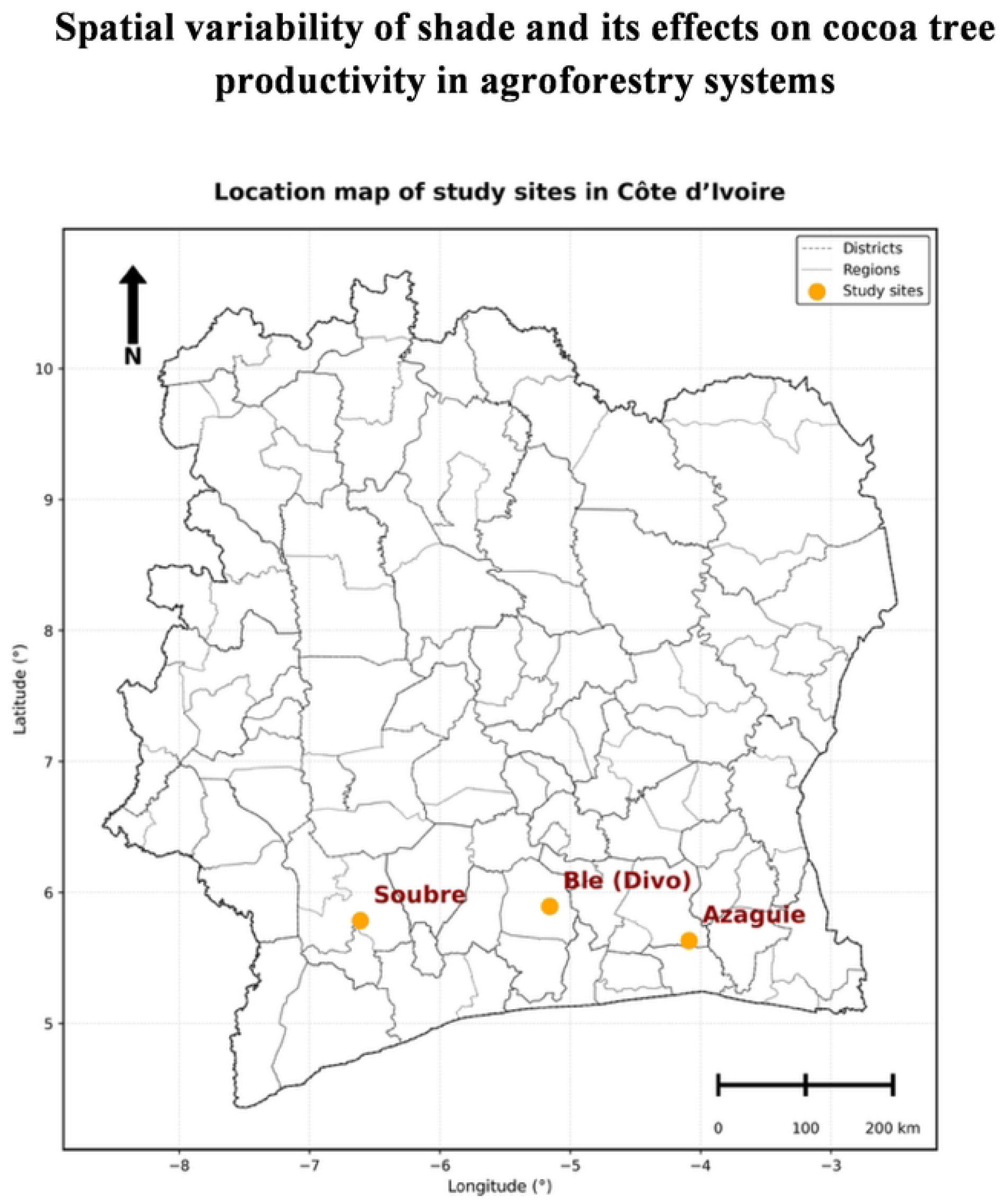
location map of study sites in Côte d’Ivoire

### 2.2. Data collection

#### 2.2.1. Shade mapping and quantification

Overstory shade was measured using an LI-191R quantum sensor on a regular grid of 25 points spaced 8 m apart, covering each 1000 m^2^ plot (Fig 2). This sampling setup was used to ensure uniform coverage of the plot and to capture fine-scale shade variability at a resolution relevant to the typical crown size of shade trees, while remaining field-practical. Photosynthetically Active Radiation (PAR) measurements inside the plot (beneath the canopy, PARi) and outside in full sun (PARe) were taken between 12:00 and 14:00 under clear skies to minimize variations due to sun angle or cloud cover [23]. The shade percentage (TO) at each grid point was calculated as:

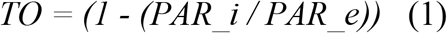

**Fig 2.**
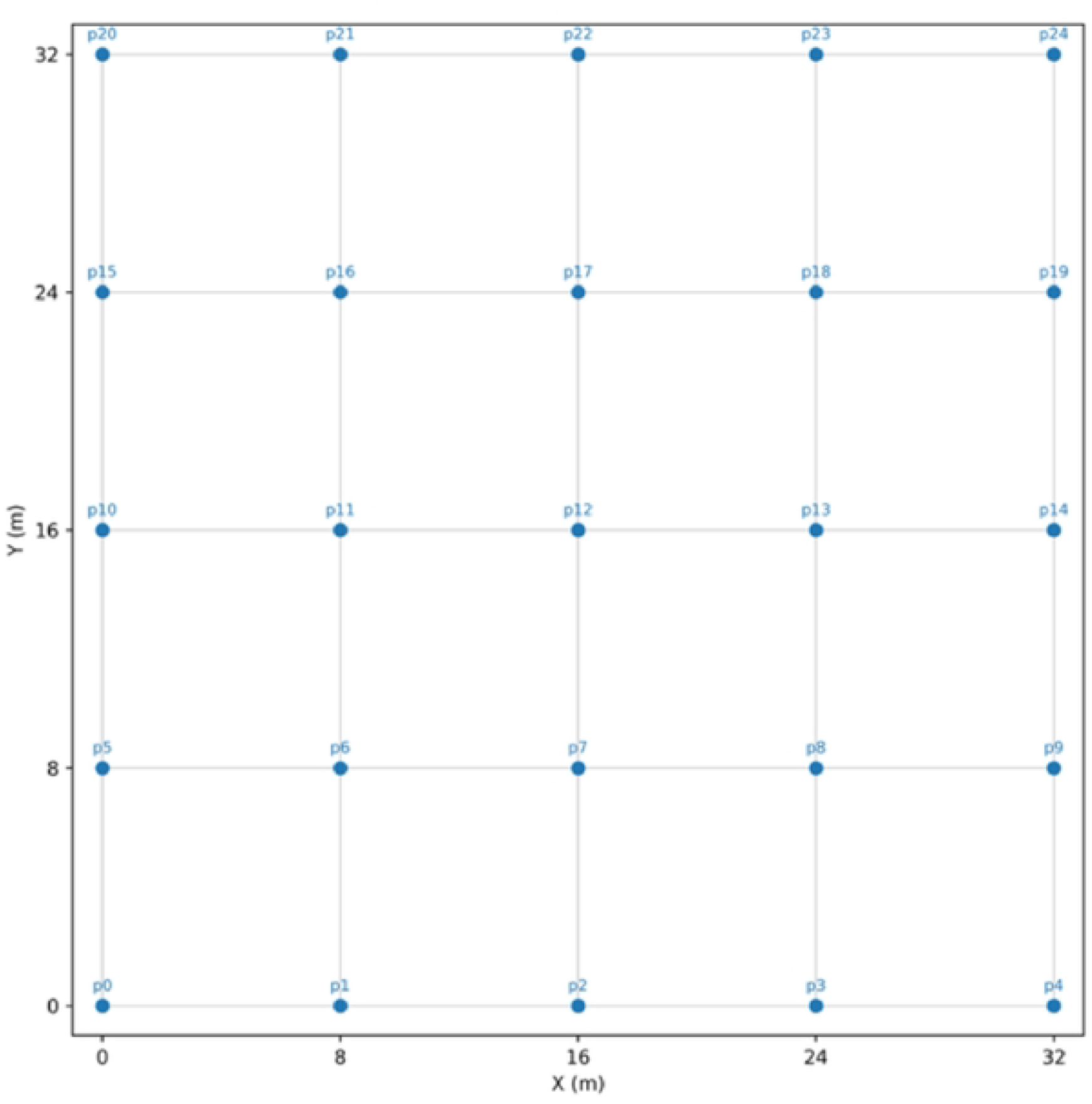
Experimental setup for measuring light radiation

This value represents the fraction of incident light intercepted by the tree canopy.

The TO values were interpolated to a continuous surface via ordinary kriging [24]. An exponential variogram model was fitted to the experimental variogram as it provided the best fit for describing the spatial dependence of our data. The grid map provides a local TO estimate (with kriging variance) at any location in the plot. All geospatial analyses were performed in Python 3.10 using the PyKrige library.

#### 2.2.2. Light transmission by shade trees

For each shade tree, we recorded its crown diameter and (x, y) coordinates relative to the plot origin. Each tree’s canopy was represented on the map by a projected circle using the measured crown diameter. A Python script then extracted the mean TO under each tree’s crown by overlaying the crown projection onto the interpolated shade map. Based on these values, each shade tree species was assigned to a light interception class: o1 (light shade, TO ≤ 30%), o2 (moderate shade, 30% < TO ≤ 60%), and o3 (dense shade, TO ≥ 60%). These thresholds distinguish low, intermediate, and high shade regimes for analysis (Table 1).

**Table 1:** Botanical diversity and light interception capacity of inventoried shade trees. o1 (light shade, ≤ 30%), o2 (moderate shade, 30–60%), o3 (dense shade, ≥ 60%)

| Taxon | Family | Capacity of light Interception |
| --- | --- | --- |
| *Alstonia boonei* | Apocynaceae | o3 |
| *Anthonotha fragrans* | Caesalpiniaceae | o2 |
| *Antiaris toxicaria* | Moraceae | o2 |
| *Canarium schweinfurtii* | Burseraceae | o2 |
| *Ceiba pentandra* | Bombacaceae | o1 |
| *Citrus muricata* | Rutaceae | o2 |
| *Citrus sinensis* | Rutaceae | o3 |
| *Cola nitida* | Malvaceae | o2 |
| *Elaeis guineensis* | Arecaceae | o1 |
| *Ficus exasperata* | Moraceae | o3 |
| *Funtumia africana* | Apocynaceae | o2 |
| *Hannoa klaineana* | Simaroubaceae | o3 |
| *Holarrhena floribunda* | Apocynaceae | o2 |
| *Mangifera indica* | Anacardiaceae | o3 |
| *Milicia excelsa* | Moraceae | o2 |
| *Morinda lucida* | Rubiaceae | o1 |
| *Morus mesozygia* | Moraceae | o3 |
| *Musanga cecropioides* | Urticaceae | o3 |
| *Parkia bicolor* | Fabaceae | o3 |
| *Persea americana* | Lauraceae | o2 |
| *Piptadeniastrum africanum* | Fabaceae | o3 |
| *Pycnanthus angolensis* | Myristicaceae | o3 |
| *Ricinodendron heudelotii* | Euphorbiaceae | o2 |
| *Spondias mombin* | Anacardiaceae | o3 |
| *Sterculia tragacantha* | Malvaceae | o3 |
| *Xylopia aethiopica* | Annonaceae | o3 |
| *Zanthoxylum gillettii* | Rutaceae | o2 |

#### 2.2.3. Cocoa productivity

The cocoa yield was evaluated by means of a manual count of pods (with a minimum length of 10 cm from base to tip) on each tree [4]. These counts were performed at the commencement of each harvest period (when pod set is representative of total yield) from early 2023 through late 2024, with two counts per year (minor and major harvest seasons).

### 2.3. Statistical analysis

Data were analyzed using geospatial and statistical approaches. First, ordinary kriging interpolation was used to model the spatial distribution of TO and generate a continuous shade map of the plot. Second, a non-parametric Kruskal-Wallis test was applied to assess the effect of shade level (full sun vs. o1, o2, o3) and shade-tree species identity on cocoa yield (pods per tree). Given that yield distributions were not normal (Shapiro-Wilk test, p < 0.05), a non-parametric test was appropriate. When the overall test was significant, Dunn’s post-hoc test with Holm’s correction (α = 0.05) was used to identify which groups differed significantly. Finally, a multiple linear regression including shade tree species (categorical predictor) and local TO (continuous predictor) was performed to quantify the specific effects of shade trees on cocoa yield. Prior to this analysis, shade tree species associated with fewer than 5 sampled cocoa trees were excluded to ensure sufficient statistical power, leaving 16 out of 27 species (approx. 86% of observations). The best-fit predictive model was selected based on Akaike’s Information Criterion (AIC). All analyses were carried out in R 4.2.3 and Python 3.10, with a significance threshold of p < 0.05.

## 3. Results

### 3.1. Shade patterns and spatial yield distribution

The kriged shade maps revealed strong spatial heterogeneity in both shade and cocoa productivity (Fig 3). Distinct high-shade zones, low-shade zones, and intermediate gradients were observed. Unshaded spots also occurred where no tree canopy was present. A total of 27 shade tree taxa were recorded in the plots. These species showed varying light interception capacities: some provided light shade (O1 ≤ 30%), others moderate shade (02, 30–60%), and others dense shade (03 ≥ 60%) (Table 1). Crown size and extent determined the area of deep shade. Regarding cocoa yield, an inverse relationship was evident between interpolated shade level and pod count: productivity tended to be higher in less shaded areas and lower in heavily shaded areas.

**Fig 3.**
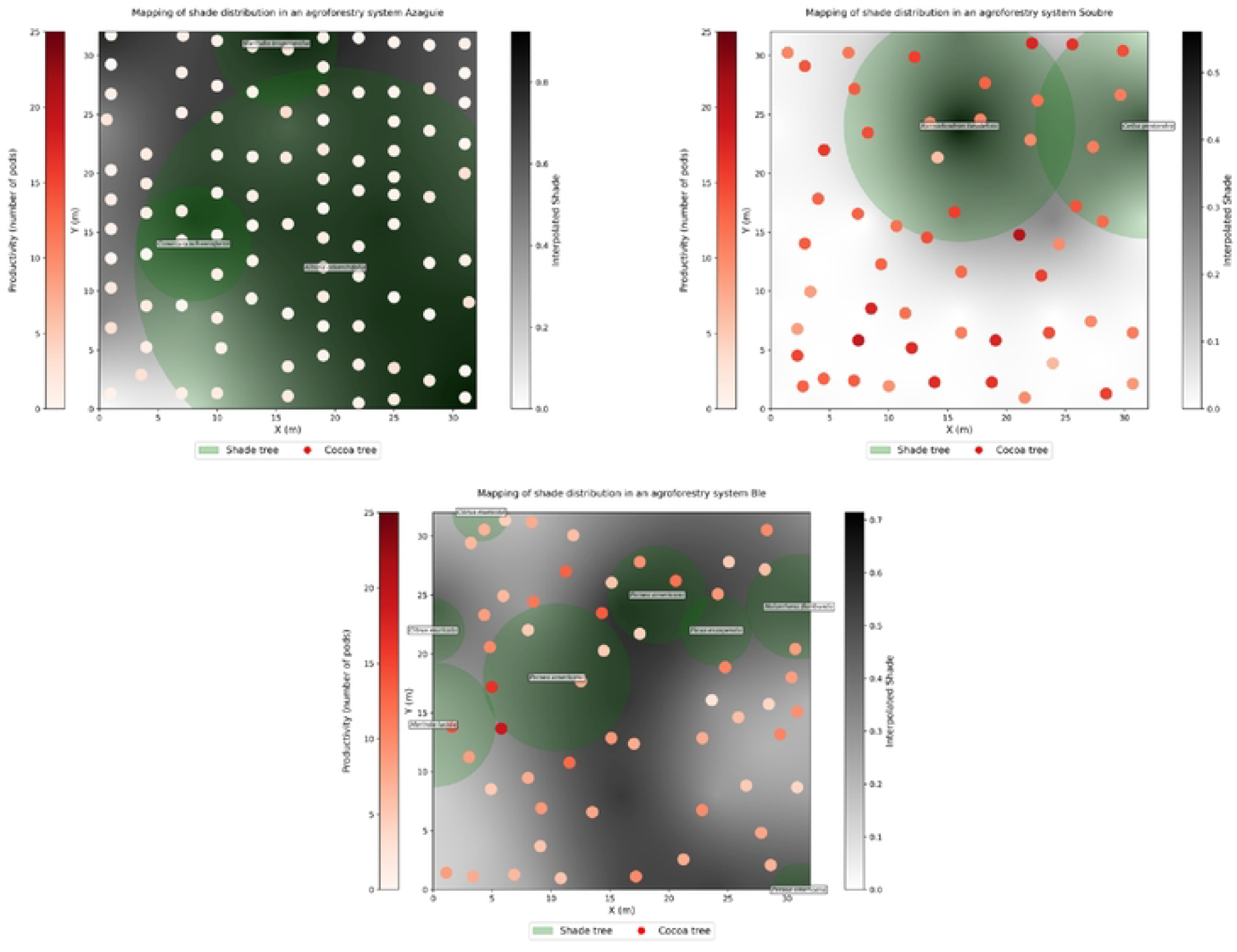
Mapping of shade fron1 three shading systems across the different study sites

### 3.2. Effect of within-plot shade heterogeneity on yield

The Kruskal-Wallis test (H = 154.1, p < 0.0001) revealed significant differences in pod yields among shade groups (Fig 4). Dunn’s post-hoc test (Holm-corrected, α = 0.05) further showed statistically distinct groupings (Table 2). The observed trend was a clear decline in yield with increasing shade. Cocoa trees in full sun produce, on average, more pods than those under any shade (12.5a). Trees under light shade (o1, ≤ 30%) averaged 9.5b pods, under moderate shade (o2, 30–60%) 2.5c pods, and under heavy shade (o3, ≥ 60%) only 2.0d pods.

**Table 2:**
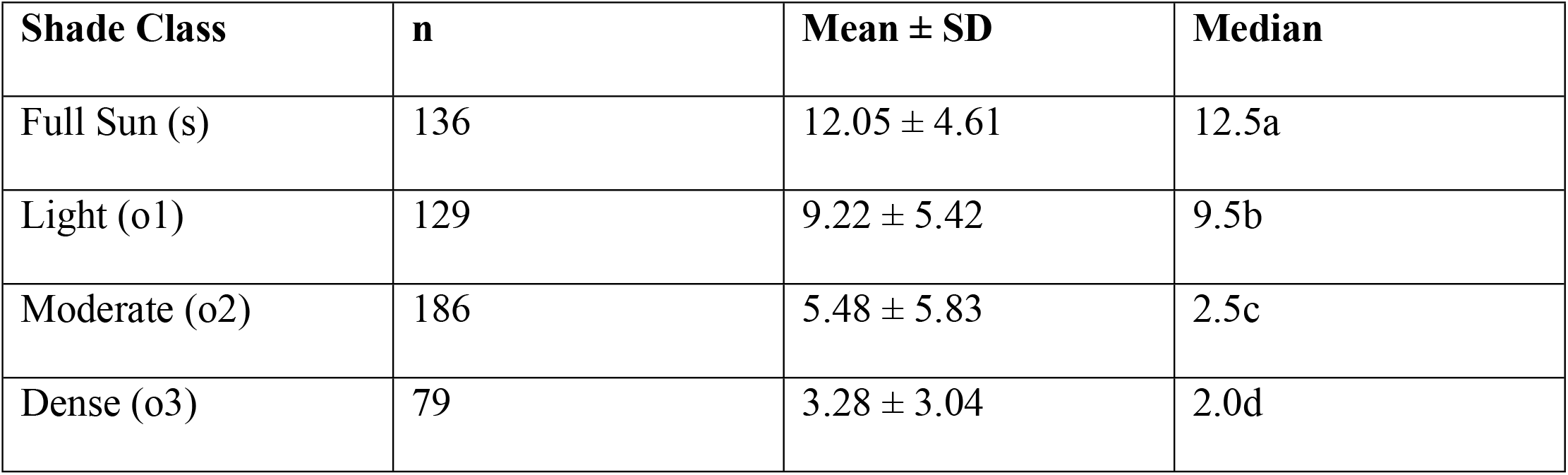
Descriptive statistics of cocoa productivity by shade class. Letters (a, b, c, d) indicate statistically distinct groups (Dunn’s post-hoc test, Holm’s correction, α = 0.05). Overall Kruskal-Wallis test: H = 154.1, p < 0.0001.X

**Fig 4.**
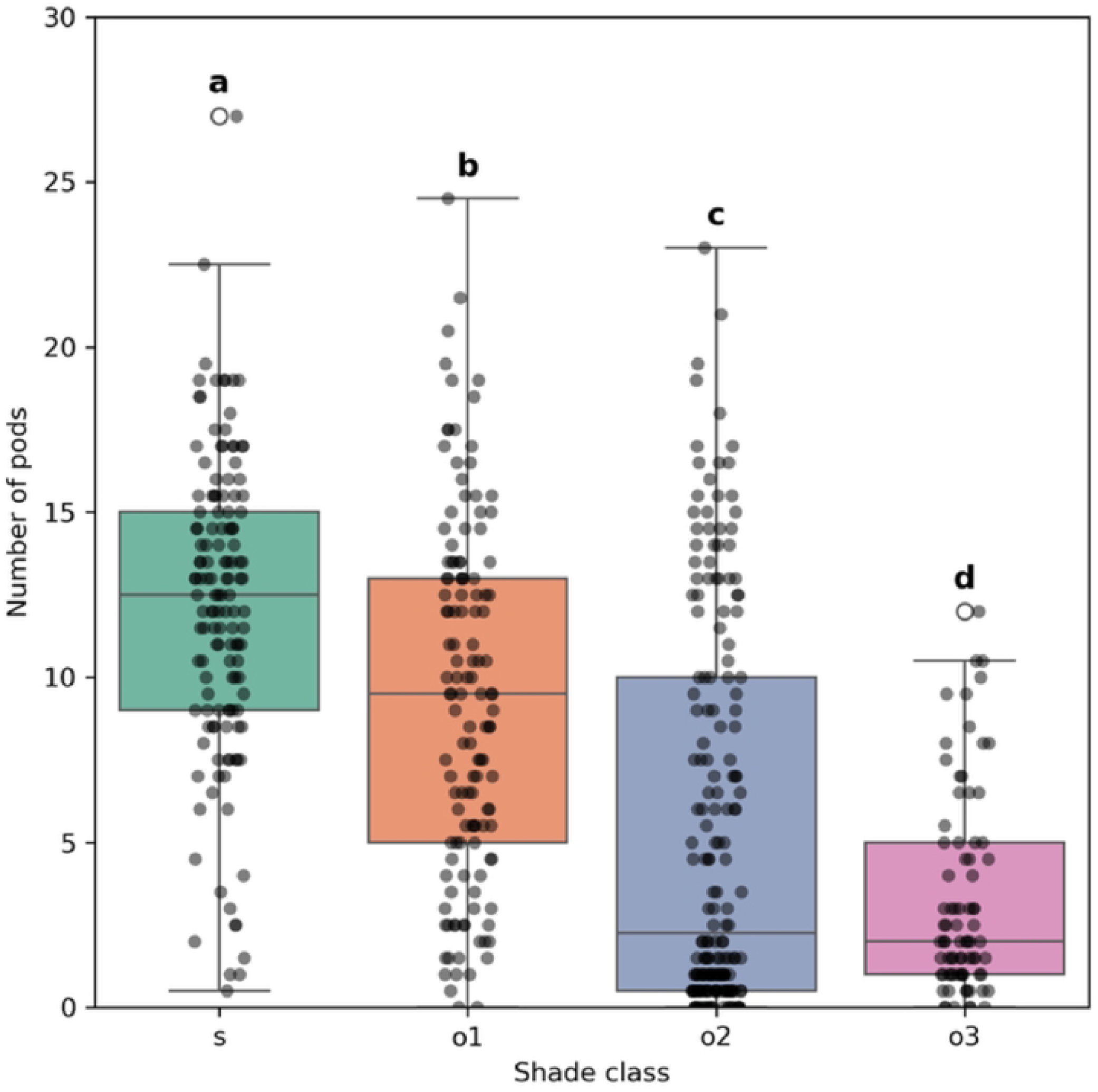
variability ofproductivity (number of pods) according to shading classes

### 3.3. Effect of specific shade-tree associations

The Kruskal-Wallis analysis also revealed significant effect of shade-tree species on cocoa yield (H = 149.7, p < 0.0001). As shown above, pod counts varied significantly with shade level and according to the associated tree species. Species providing the least shade such as *Ceiba pentandra* and *Elaeis guineensis* (TO ≤ 30%) were associated with the highest cocoa yields. In contrast, species causing heavy shade such as *Mangifera indica* were associated with the lowest yields.

For the predictive analysis, we first filtered out species with < 5 observations, retaining 16 species (86.2% of the data). The optimal regression model (including shade tree species and their light interception capacity) explained approximately 68.4% of the variance in cocoa yield (R^2^ = 0.684, AIC = 854.7) (Fig 5). The shade tree species associated with the greatest cocoa yields were *Ceiba pentandra* (median = 13.5 pods/tree), *Elaeis guineensis* (11.8), *Persea americana* (9.5), and *Ricinodendron heudelotii* (7.8) (Fig 6).

**Fig 5.**
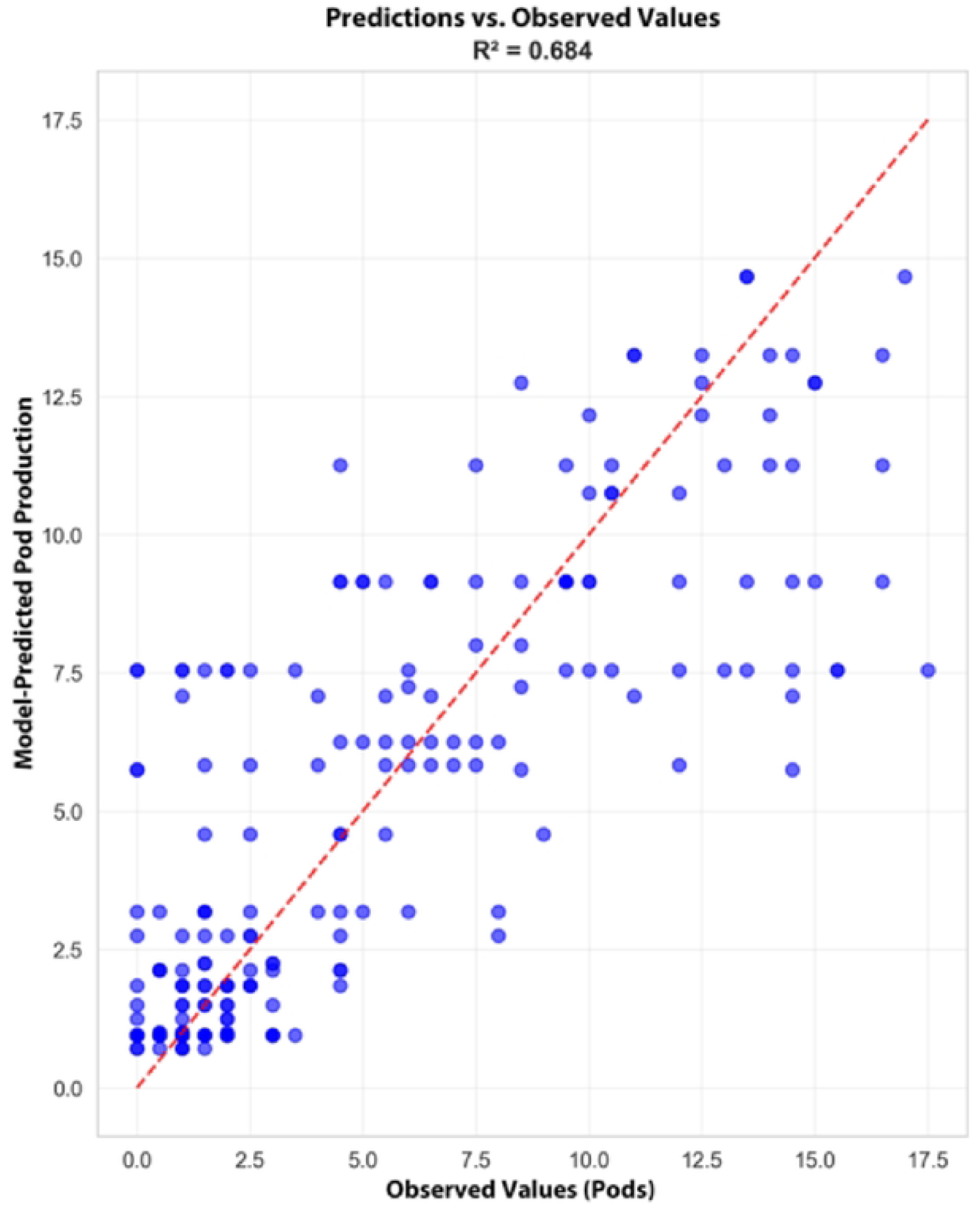
Performance of the predictive 1nodel on the influence of shade trees on cocoa productivity

**Fig 6.**
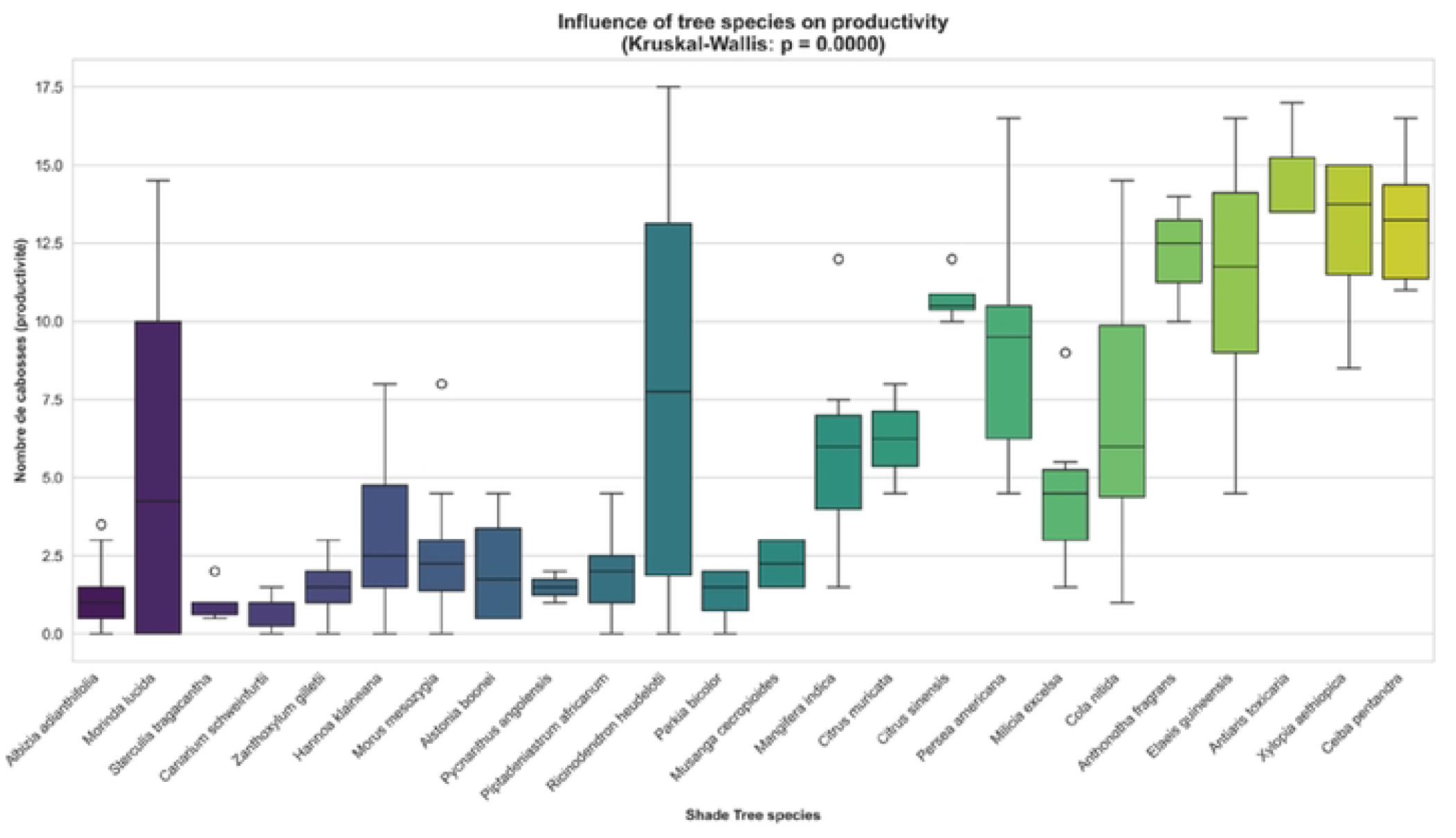
Distribution of productivity based on shade tree and cocoa tree associations

## 4. Discussion

The spatial distribution of shade in cocoa agroforestry, as revealed by our geostatistical analysis, exhibits strong heterogeneity due to the phenological and morphological diversity of companion trees [14,25,26]. This heterogeneity, or multi-shade, is a key lever for creating a favorable microclimate tempering abiotic stresses like heat and drought while influencing pest and disease dynamics [11–13,15,27]. Our results indicate that 68.4% of the variance in cocoa yield is explained by the combination of shade tree species and local shade level, confirming that interactions between companion trees and cocoa trees are crucial, especially when proximity amplifies shade effects [17].

Cocoa yields drop considerably beyond 60% shade, corroborating observations by [16,28] that excessive shade reduces photosynthesis and limits yield [29,30]. Conversely, moderate shade around 30% appears most conducive, striking a balance between climate regulation and light availability, in line with [6] and [9].

Our findings also confirm the species-specific nature of shade cocoa interactions *Ceiba pentandra* clearly stands out for its light, consistent shade that favors productivity a result already documented with respect to its beneficial litter quality and deep rooting that reduce competition for water. Other species like *Elaeis guineensis* and *Persea americana* also prove advantageous; their moderate shade helps maintain a balance between light and protection, which explains their frequent use in some cocoa-growing regions of Ghana.

In contrast, species such as *Mangifera indica* appear detrimental to production. While its very dense shade, which can reduce light transmission to as low as 3%, is the primary cause of yield loss, other competitive mechanisms may also be at play. For instance, *Mangifera indica* is known for its extensive, shallow root system that could create intense below-ground competition for water and nutrients, or potential allelopathic effects could further inhibit cocoa growth. These alternative hypotheses warrant further investigation [9,27].

Notably, while reducing shade enhances cocoa productivity, denser shade can provide other ecosystem services, for instance by limiting certain heat-sensitive fungal diseases, supporting biodiversity [31], and contributing to carbon sequestration and long-term resilience [32]. The sharp yield decline above 60% shade may not only be due to light limitation for photosynthesis [29,30,33] but could also be exacerbated by higher humidity under dense canopies, which favors pathogens. This highlights a critical trade-off: optimizing cocoa agroforestry cannot focus solely on maximum yield but must pursue a balance between production and sustainability. Our results suggest that a “shade mosaic” design could be a promising management strategy, creating zones of optimal light (~30%) for production interspersed with denser shade patches that serve as biodiversity refuges and microclimate buffers.

The originality of our study lies precisely in this geospatial analysis of intra-plot multi-shade. By showing that spatial shade variability is a major explanatory factor for yield differences among cocoa trees in the same plot, our work fills a gap highlighted by many authors [16–18] and offers concrete prospects for cocoa agroforestry adapted to contemporary climate and production challenges.

## 5. Conclusion

The heterogeneous spatial distribution of shade, illustrated by ordinary kriging analysis, plays a crucial role in cocoa productivity. The morphological and architectural diversity of companion trees creates shade gradients across the agroforestry plot, which modulate the microclimate and significantly influence cocoa yields. A low shade of around 30% has been identified as being optimal for high cocoa productivity, underscoring the significance of selecting shade tree species that are able to transmit low to moderate levels of light. Among the species studied, *Ceiba pentandra* is notable for its favourable characteristics in cocoa agroforestry, specifically its capacity to provide gentle shade and its positive effects on soil. It has been demonstrated that other species, such as *Elaeis guineensis* and *Persea americana*, which provide light to moderate shade, are also beneficial for cocoa. However, it has been shown that species casting overly dense shade, such for example *Mangifera indica*, can become counterproductive. The present study provides robust scientific bases for the implementation and design of cocoa-based agroforestry systems. However, these conclusions should be further explored through long-term monitoring and a detailed understanding of the functional traits that govern light interception by companion trees.

## Acknowledgments

This study was conducted within the framework of the Cocoa4Future (C4F) project, which is funded by the European DeSIRA Initiative under grant agreement No. FOOD/2019/412-132 and by the French Development Agency.

## 6. Data Availability Statement

All relevant data are within the manuscript and its Supporting Information files.

## Spatial variability of shade and its effects on cocoa tree productivity in agroforestry systems

